# Sex and origin-specific inbreeding effects on flower attractiveness to specialised pollinators

**DOI:** 10.1101/2021.01.08.425842

**Authors:** Karin Schrieber, Sarah Catherine Paul, Levke Valena Höche, Andrea Cecilia Salas, Rabi Didszun, Jakob Mößnang, Caroline Müller, Alexandra Erfmeier, Elisabeth Johanna Eilers

## Abstract

We investigate whether inbreeding has particularly fatal consequences for dioecious plants by diminishing their floral attractiveness and the associated pollinator visitation rates disproportionally in females. We also test whether the magnitude of such effects depends on the evolutionary histories of plant populations. We recorded spatial, olfactory, colour and rewarding flower attractiveness traits as well as pollinator visitation rates in experimentally inbred and outbred, male and female *Silene latifolia* plants from European and North American populations differing in their evolutionary histories. We found that inbreeding specifically impairs spatial and olfactory attractiveness. Our results support that sex-specific selection and gene expression partially magnified these inbreeding costs for females, and that divergent evolutionary histories altered the genetic architecture underlying inbreeding effects across population origins. Moreover, they highlight that inbreeding effects on olfactory attractiveness have a huge potential to disrupt interactions among plants and specialist moth pollinators, which are mediated by elaborate chemical communication.

## 1. Introduction

Plant-pollinator interactions are of central importance for the emergence and maintenance of global biodiversity (Crepet and Niklas, 2009; Ollerton, 2017) and provide ecosystem services with tangible economic and cultural value (Gill et al., 2016; Porto et al., 2020). Global change continues to disrupt these interactions by altering the physiology, phenology and particularly the spatial distribution of component species (Burkle et al., 2013; Vanbergen and Initiative, 2013; Glenny et al., 2018). Habitat degradation and fragmentation reduce the size and connectivity of plant populations, which results in lowered pollinator visitation rates (Aguilar et al., 2006; Dauber et al., 2010). Population retraction and isolation may also affect plant attractiveness to pollinators at the individual level by increasing inbreeding rates (Carr et al., 2014). However, mechanistic insight into the consequences of inbreeding for plant traits attracting pollinators and feedbacks on visitation rates are to date scarce. Likewise, it is largely unexplored whether inbreeding effects on attractiveness can be magnified by sex-specific selection and gene expression in dioecious, pollinator-dependent plants and whether populations can escape such constraints by purging of the genetic load.

Inbreeding raises homozygosity in the offspring generation. This can enhance the phenotypic expression of deleterious recessive mutations (i.e., dominance) and reduce heterozygote advantage (i.e., over-dominance) with fatal consequences for Darwinian fitness (i.e., inbreeding depression) (Charlesworth and Willis, 2009). Plant inbreeding may also disrupt interactions with pollinators by directly reducing floral attractiveness. Floral attractiveness is a complex syndrome composed of i) the scent bouquet as determined by the composition of floral volatile organic compounds (VOC) such as terpenoids, benzenoids and phenylpropanoids (Muhlemann et al., 2014; Borghi et al., 2017); ii) spatial traits including the number, size, shape, and orientation of flowers (Dafni et al., 1997); iii) flower colour as defined by the composition of pigments with wavelength-selective light absorption and the backscattering of light by petal surface structures (van der Kooi Casper J. et al., 2016; Borghi et al., 2017); and iv) the quality and quantity of rewards such as nectar, pollen, oviposition sites or shelter (Simpson and Neff, 1981). These cues are particularly efficient in attracting pollinators across either long (scent), medium (spatial traits, colour) or short (rewards) distances (Dafni et al., 1997; Muhlemann et al., 2014) and act synergistically in determining visitation rates. Although in a few cases inbreeding has been shown to alter single components of flower attractiveness (Ivey and Carr, 2005; Ferrari et al., 2006; Haber et al., 2019), insight into syndrome-wide effects is lacking.

The magnitude and slope of inbreeding effects in plants can vary across environments, since local conditions partly determine the selective value of recessive alleles unmasked by inbreeding (Fox and Reed, 2011). While the influence of environmental stress on the expression of inbreeding depression is well studied, the effects of plant sex, which considerably shapes an individuals’ interaction with its environment, remain largely unexplored. Individuals of dioecious plant species invest into either male or female reproductive function. This partitioning goes along with different life histories, resource demands, stress susceptibilities and, consequently sex-specific selection regimes in identical habitats (Moore and Pannell, 2011; Barrett and Hough, 2013). Sex-specific selection may modify the magnitude of inbreeding depression in dioecious plants. Studies on animals reported higher inbreeding depression in female than male individuals resulting from higher reproductive investment and prolonged life cycles in the former (Ebel and Phillips, 2016). In plants, such relations have rarely been investigated (Teixeira et al., 2009), and if so, not with a focus on floral attractiveness traits. Compared to males, female plants invest typically less in attraction of pollinators, since their reproductive success is more limited by the availability of resources for fruit production than by the availability of mates (Moore and Pannell, 2011; Barrett and Hough, 2013). If inbreeding effects on attractiveness are more detrimental for females at the same time, the relative frequency of pollinator visits may be biased towards male plants, with devastating consequences for the effective size and growth of populations.

Plant populations may escape progressive retractions under increased inbreeding rates by purging. Inbreeding unmasks deleterious recessive mutations, which facilitates their selective removal from the population gene pool and may result in a rebound of fitness when the demographic bottleneck is intermediate (Crnokrak and Barrett, 2002). Plant species that have successfully colonised distant geographic regions provide perfect models for studying the relevance of purging in natural plant populations. As colonization events are associated with successive demographic bottlenecks, purging is expected to be one determinant for the successful establishment and proliferation of plant populations in novel habitats (Facon et al., 2011; Schrieber and Lachmuth, 2017). However, only few empirical studies verified the role of purging in plant colonization success by revealing significantly lower inbreeding depression following experimental crossings in invasive than native plant populations (Rosche et al., 2016; Schrieber et al., 2019). Again, the focus of these studies was on fitness components rather than floral attractiveness traits. Yet floral attractiveness is a key for successful colonization in pollinator-dependent species introduced into novel communities (Morales and Traveset, 2009), especially if plants are not capable of selfing (e.g., dioecious). Attractiveness syndromes are likely under strong selection in such species, which should rapidly purge deleterious recessive mutations affecting spatial, scent, colour and rewarding flower traits.

In the present study, we investigated inbreeding effects on plant attractiveness to pollinating insects and its dependence on sex and population origin using the plant species *Silene latifolia* Pior. (Caryophyllaceae) and its crepuscular moth pollinators. Natural *S. latifolia* populations partly suffer from biparental inbreeding due to limited seed and pollen dispersal (McCauley, 1997). As inbreeding reduces not only fitness (Teixeira et al., 2009), but also impairs interactions with herbivorous insects in *S. latifolia* (Schrieber et al., 2018, 2019), a disruption of plant-pollinator interactions can be expected for this species. Moreover, the dioecious reproductive system of *S. latifolia* provides the opportunity to quantify variation in the magnitude of inbreeding effects in these traits among females and males. Finally, the species expanded successfully from parts of its native distribution range in Europe to North America in the early 19th century (Keller et al., 2009, 2012), which may have given rise to purging events. We assessed flower attractiveness *via* spatial traits, headspace scent composition, colour and rewards and quantified pollinator visitation in experimentally inbred and outbred male and female *S. latifolia* individuals from European and North American populations. We hypothesised that: i) inbreeding reduces plant attractiveness and this effect is more pronounced ii) in female than male plants, and iii) in European than North American populations. iv) The combined effects of inbreeding, sex and population origin cause feedback on pollinator visitation rates.

## 2. Material & Methods

### 2.1 Study species

*Silene latifolia* shows a distinct moth pollination syndrome with large, white and funnel-shaped flowers (Dafni et al., 1997). The flowers open from dusk till mid-morning to release a scent bouquet composed of more than 60 VOC, whereby emission peaks around dusk (Dötterl et al., 2005, 2009; Mamadalieva et al., 2014). A substantial fraction of VOC in this blend triggers antennal and behavioural responses in nocturnal moths (Dötterl et al., 2006). Nectar production peaks three to four days after flower opening and is just as floral scent emission reduced after pollination (Gehring et al., 2004; Dötterl et al., 2005; Muhlemann et al., 2006). *Silene latifolia* exhibits various sexual dimorphisms with male plants producing more and smaller flowers that excrete lower volumes of nectar with higher sugar concentrations as compared to females (Gehring et al., 2004; Delph et al., 2010) but no clear sex-specific patterns in floral scent composition (Dötterl & Jürgens 2005 but see Waelti *et al*. 2009).

Various diurnal generalist pollinators as well as crepuscular moths visit *S. latifolia* flowers. The latter, including the specialist *Hadena bicruris* Hufn. (Lepidoptera: Noctuidae), were shown to be the most efficient pollinators for *S. latifolia* (Young, 2002). *Silene latifolia* and *H. bicruris* form a well-studied nursery pollination system, in which female moths pollinate female plants while ovipositing on the flower ovaries to provide their larvae with developing seeds. Pollination services provided by male *H. biruris* over-compensate the costs of seed predation by their offspring (Labouche and Bernasconi, 2010). The activity of *H. bicruris* peaks at dusk between May and July (Bopp and Gottsberger, 2004). *Hadena bicruris* is abundant in 90 % of European *S. latifolia* populations but has not yet been introduced to North America. Other nocturnal moths such as including *Hadena ectypa* Morrison (Lepidoptera: Noctuidae) provide main pollination services to *S. latifolia* in the invaded range without imposing costs by seed predation (Young, 2002; Castillo et al., 2014).

### 2.2 Plant material

We collected seed capsules from five female individuals (maternal families) in each of eight European and eight invasive North American *S. latifolia* populations (Fig. S1, Table S2). Seeds from all maternal families were germinated and plants were grown under controlled greenhouse conditions for experimental crossings within populations. Each female individual from the P-generation received pollen from a male derived from the same maternal family (inbreeding) and pollen from a male derived from a different maternal family within the same population (outcrossing) at separate flowers (Fig. S3). The field sampling, rearing conditions and experimental crossing are described in detail in (Schrieber et al., 2018, 2019). Seeds were dried and stored at room temperature until use.

For the experiment, we grew plants from the F1 generation under greenhouse conditions (16: 8 hr light: dark at 20 / 10 °C ± 6 °C). After the onset of flowering, we randomly chose one female and one male individual per breeding treatment (inbred, outbred) × maternal family (1-5) × population (1-8) × origin (Europe, North America) combination, resulting in 320 plant individuals for the experiment. Using these individuals, we assessed the combined effects of breeding treatment, plant sex and population origin on flower traits mediating plant attractiveness to pollinators and pollinator visitation rates during the summers 2019 and 2020. Plants were grown in 3 L (2019) and 6 L (2020) pots filled with a 3 : 1 mixture of potting soil (TKS2 Instant Plus, Floragrad, GE) and pine bark (Pine Bark 1-7 mm, Oosterbeek Humus Producten, NL). They were kept in pots with randomised positions either in the greenhouse, a common garden with sealed ground (54.346794°N, 10.107990°E, 19 m elevation) or a field site covered by an extensively used meadow (54.347742°N, 10.107661°E, 19 m elevation) for different parts of data acquisition. For an overview of the time schedule, locations and exact sample sizes for data acquisition see Table 1. Plants received water, fertilization (UniversolGelb 12-30-12, Everris-Headquarters, NL) or treatment with biological pest control agents (Katz Biotech GmbH, GE) when necessary for the entire experimental period.

**Table 1:** Overview of locations, times and sample sizes for data acquisition.

| Trait category | Location | Acquisition time<br>(year, month,<br>duration) | Nesting and intended total<br>sample size | Realised<br>total sample<br>size | Replicates per<br>group (breeding<br>treatment × sex ×<br>origin combination) | Reason for sample size<br>reduction |
| --- | --- | --- | --- | --- | --- | --- |
| Spatial flower traits<br>(plant height,<br>flower number) | Greenhouse &<br>common garden | 2019<br>Jun., Jul., Aug.<br>5 d respectively | 2 breeding treatments × 2 sexes ×<br>5 maternal families × 8<br>populations × 2 origins = 320 | 316 | 36-40 | 4 individuals died |
| Flower scent | Greenhouse | 2019<br>Jul.<br>8 hours | 2 breeding treatments × 2 sexes ×<br>3 maternal families × 8<br>populations × 2 origins = 192 | 192 | 23-35 | - |
| Flower colour | Common garden | 2019<br>Aug.<br>2 weeks | 2 breeding treatments × 2 sexes ×<br>5 maternal families × 8<br>populations × 2 origins = 320 | 286 | 23-25 | 4 individuals died, no<br>flowers available for<br>remaining plants |
| Spatial flower traits<br>(petal limb area<br>and expansion) | Common garden | 2019<br>Aug.<br>2 weeks | 2 breeding treatments × 2 sexes ×<br>5 maternal families × 8<br>populations × 2 origins = 320 | 286 | 23-35 | 4 individuals died, no<br>flowers available for<br>remaining plants |
| Pollinator visitation<br>rates | Field site | 2020<br>May-Jul.<br>8 weeks | 2 breeding treatments × 2 sexes ×<br>5 maternal families × 8<br>populations × 2 origins = 320 | 316 | 36-40 | 4 individuals died |
| Floral rewards | Common Garden | 2020<br>Aug.<br>4 weeks | 2 breeding treatments × 2 sexes ×<br>5 maternal families × 8<br>populations × 2 origins = 320 | 280 | 30-40 | 4 individuals died, no<br>flowers available for<br>remaining plants |

### 2.3 Plant attractiveness

#### 2.3.1 Spatial flower traits

We determined the inflorescence height and the number of fully opened flowers per individual. These traits were acquired thrice in intervals of two and six weeks to account for phenological variation, and we refer to their average values hereinafter. The size of *S. latifolia* flowers was not assessed *via* the length of their petal limbs as in previous studies, since this estimate does not account for the severe variation in their overall shape (Fig. S4). Instead, we assessed the exact area covered by all petal limbs and the expansion of the corolla (i.e., the area covered by the smallest possible circle drawn around all five petal limbs). Both traits were derived from digital images taken from one well-developed and fully-opened flower per plant (see 2.3.3 for further details) using the software ImageJ 1.47t (Rueden et al., 2017).

#### 2.3.2 Flower scent

For characterization of flower scent, we trapped the headspace VOC of *S. latifolia* flowers on absorbent polydimethylsiloxane (PDMS) tubing following the method of Kallenbach et al. (2014, 2015). We placed the plants in a spatial distance of 50 cm to one another in the greenhouse and maintained high air ventilation one week prior to and during VOC collection. We selected one well-developed flower per individual and enclosed it in a VOC collection unit (Fig. S5). The collection units consisted of polypropylene cups with lids (50 mL, Premium Line, Tedeco-Gizeh, GE), both having holes (diameter 15 mm) to prevent heat and waterlogging. They were fixed *via* wooden sticks at the exterior of the plant pot. In addition, 14 control collection units were fixed on empty plant pots and positioned throughout the greenhouse. Prior use, the absorbent PDMS tubes (length 5 mm, external diameter 1.8 mm, internal diameter 1 mm; Carl Roth, GE), were cleaned with solvents and heat as described in Kallenbach et al. (2014). Two PDMS tubes were added to each collection unit and remained in the floral headspace between 9 pm and 5 pm, which is the time of peak scent emission in *S. latifolia* (Dötterl et al., 2005). Afterwards, the PDMS tubes were removed and stored at −20 °C in sealed glass vials until analysis *via* thermal desorption-gas chromatography-mass spectrometry (TD-GC-MS, TD 30 - GC 2010plus - MS QP2020, Shimadzu, JP).

Trapped VOC were desorbed from PDMS tubes for 8 min at 230 °C under a helium flow of 60 mL min^−1^ and adsorbed on a Tenax^®^ cryo-trap with a temperature of −20 °C. From the trap, compounds were desorbed at 250 °C for 3 min, injected to the GC in a 3:10 split mode, and migrated with a helium flow of 1.6 mL min^−1^ on a VF5-MS column (30 m × 0.25 mm + 10 m guard column, Agilent Technologies, USA). The GC temperature program started at 40 °C for 5 min and increased to 125 °C at a rate of 10 °C min^−1^ with a hold time of 5 min and to 280 °C at a rate of 30 °C min^−1^ with a hold time of 1 min. Line spectra (30 - 400 m/z) of separated compounds were acquired in quadrupole MS mode. An alkane standard mix (C8-C20, Sigma Aldrich, GE) was analysed under the same conditions in order to calculate Kovats retention indices (KI) for targeted compounds (Kováts, 1958).

Compounds were identified by comparing the KI and mass spectra with those of synthetic reference compounds, where available, and with library entries of the National Institute of Standards and Technology (NIST) (Smith et al., 2004), Pherobase (El-Sayed, 2011), the PubChem database (Kim et al., 2016), and (Adams, 2007). Compounds were not quantified but the relative intensity of the total ion chromatogram of peaks was compared. Control samples (collection units without flowers), and blanks (cleaned PDMS tubes) were used to identify and exclude contaminations, leaving a total number of 70 VOC (Table S5). For targeted statistical analyses, we focused on those VOC that evidently mediate communication with *H. bicruris*. We analysed the Shannon diversity per plant (calculated with R-package: vegan v.2.5-5, Oksanen et al. 2019) for VOC that elicit electrophysiological responses in the antennae of *H. bicruris*, as well as the relative intensity of three lilac aldehyde isomers, which trigger oriented flight and landing most efficiently in *H. bicruris* according to Dötterl et al. (2006) (Table S5).

#### 2.3.3 Flower colour

Flower colour was quantified using a digital image transformation approach that accounts for the visual system of the pollinator as well as natural light conditions (Troscianko & Stevens 2015). Images were acquired in the common garden after plants had acclimated to ambient light conditions for three weeks. All images were taken during one hour of dusk time on rain-free days in order to fit the natural light conditions perceived by *H. bicruris* (Bopp and Gottsberger, 2004). We picked one well developed, fully opened flower per plant and inserted it into a black ethylene vinyl acetate platform equipped with two reflectance standards (PTFB 10 %; Spectralon 99 % Labsphere, Congleton, UK) and a size standard (Fig. S7). The platform had a fixed location in the field and was oriented towards the setting sun. Raw images were taken with a digital camera (Samsung NX1000, JP) converted to full spectrum sensitivity (300-1000 nm) *via* removal of the sensor’s filter and fitted with a UV sensitive lens (Nikon EL 80-mm, JP). We took images in the visible and in the UV part of the light spectrum by fitting an UV and infrared (IR) blocking filter (UV/IR Cut, transmittance 400 - 700 nm, Baader Planetarium, GE) and an UV pass plus IR block filter (U-filter, transmittance 300 - 400 nm, Baader Planetraium, GE) to the lens, respectively. All images were taken as RAWs with an aperture of 5.6, an iso of 800 and a shutter speed varying according to light conditions.

Images were processed using the Multispectral Image Calibration and Analysis (MICA)-Toolbox plugin (Troscianko and Stevens, 2015) in Image J 1.47t (Rueden et al., 2017). They were linearised to correct for the non-linear response of the camera to light intensity and equalised with respect to the two light standards in order to account for variation in natural light perceived among images (Stevens et al., 2007). All petals were selected for analysis, and the reproductive organs and para-corolla were omitted. Linearised images were then mapped to the visual system of a nocturnal moth. As the visual system of *H. bicruris* is unexplored, we used the tri-chromatic visual system of *Deilephila elpenor* L. (Lepidoptera: Sphingidae), which includes three rhodopsins with absorption maxima of 350 nm (UV), 440 nm (blue), and 525 nm (green) (Johnsen et al., 2006). We considered this system to be comparable to that of *H. bicruris*, given the similar activity behaviour of adults, morphological similarity of the preferred plant species (*Lonicera periclymenum* L. (Caprifoliaceae) with white-creamy funnel-shaped flowers) and overlapping distribution ranges. We fitted the images to a cone catch model incorporating i) the spectral sensitivity of our Samsung NX1000-Nikkor EL 80mm 300-700 nm camera (data derived from Troscianko & Stevens 2015), ii) the spectral sensitivities of the three photoreceptors in the *D. elpenor* compound eye (data derived from Johnsen et al. 2006); and iii) the spectral composition of sun light during dusk (data derived from Johnsen et al. 2006).

#### 2.3.4 Floral rewards

As moths forage on liquids only, we measured nectar as floral reward. We selected one well-developed, closed flower bud per plant in the common garden and enclosed it in a transparent mesh bag (Organza mesh bags, Saketos, GE) until harvest to avoid pollination and nectar removal. All flowers were harvested at noon of the fourth day after opening and were stored immediately at 4 °C until processing to prevent further nectar secretion. Nectar was extracted into 1-2 μL microcapillary tubes (Minicaps NA-HEP, Hirschmann Laborgeräte, GE). The length of the nectar column was measured with callipers to determine the exact volume. Nectar sugar content was analysed with a refractometer adjusted for small sample sizes (Eclipse Low Volume 0-50° brix, Bellingham and Stanley, UK). Since nectar volume trades off against nectar quality in pollinator attraction (Cnaani et al., 2006), we addressed floral rewards in *S. latifolia via* the total amount of sugar excreted per flower as calculated based on the following equation: 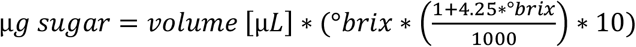.

### 2.4 Pollinator visitation rates

We quantified visits by crepuscular pollinators belonging to the order of Lepidoptera at the field site. For this purpose, plants were arranged in plots (1.5 × 1.5 m, distance among plants = 0.5 m) that consisted of eight individuals representing all populations from one breeding treatment × sex × origin combination. Each of the possible combinations (N = 8) was replicated five times at the level of maternal families, resulting in a total number of 40 plots (N = 320 plants in total). Plots were spaced from each other at a distance of 6 m in order to provide pollinators with the choice of visiting specific breeding treatment × sex × origin combinations (Glenny et al., 2018). The position of plots and plants within plots was fully randomised (Fig. S8). We performed 14 observation trials between May and July to cover the annual peak activity of *H. bicruris* (Bopp and Gottsberger, 2004). Each trial comprised five minutes observation time for each of the plots (total observation time: 2800 min, observation time per plot: 70 min) and was completed within one hour in the dawn time by four observers. The exact daytime of observation was acquired at the plot-level for each of the trials. Plant and flower visits were determined at the plant individual level. If a moth had first contact with a flower, this was counted as a plant visit. The number of approached flowers per plant during a visit was counted until a moth either left or switched to another plant. The number of plant and flower visits per trial were averaged at the plot level for further analyses. The number of visiting moth individuals and moth species were not determined. The vast majority of visits were performed by *H. bicruris* (personal observation).

Seed set was not quantified, since the exchange of pollen within and among breeding treatments and population origins could not be controlled, which would have severely confounded their effects. However, we consistently removed ripe fruits (brownish colour, dry fruit wall showing first signs of dehiscence) from female plants in order to prevent the escape of genetic material from our experiment without impeding resource investment in fruit and seed production.

### 2.5 Statistical analyses

All statistical analyses were performed in R v4.0.3 (R Development Core Team, 2020) with (generalised) linear mixed-effects models (LMMs: R-package lme4 v1.1-23, Bates et al. 2014), GLMMs: R-package glmmTMB v1.0.2.1, Brooks et al. 2017). Models for responses reflecting spatial traits, floral scent, colour and rewards included the predictors breeding treatment, sex and origin as well as all possible interactions among these factors. The latitudinal coordinate of the population origin was included as covariate in all models, whereas the exact age of the plant individuals (accounts for difference of 12 d in planting date) was included only in models for flower scent, which was acquired in early phases of the experiment. Both covariates were centred and scaled (i.e., subtraction of mean and division by standard deviation). The random effects for plant attractiveness models were population, affiliation of paternal plant in P-generation to field collected family nested within population, and affiliation of maternal plant in P-generation to field collected family nested within population. Models for pollinator visitation rates included the predictors breeding treatment, sex and origin as well as all possible interactions among them, the covariate daytime (centred and scaled), and the random effects of plot and trail (latitude of population origin, population, maternal and paternal affiliation not included, since data were averaged on plot level, see 2.4). Several of the described models included count data responses with an access of zeroes (intensity of lilac aldehydes and pollinator visitation rates). These models were additionally fitted with zero-inflation formulas. The fit of lilac aldehydes models was best when including only an intercept model for zero-inflation, whereas the fit of pollinator visitation rate models was best when including the same predictors and random effects in the conditional and zero inflation part of the model.

All of the described models (Table 1) were validated based on checking plots (quantile-quantile, residual *versus* fitted) and tests provided in the R-package DHARMa v0.3.3.0 (Hartig, 2020). Sum-to-zero contrasts were set on all factors for the calculation of type III ANOVA tables based on Wald χ^2^ tests (R-package: car v3.0-10, Fox & Weisberg 2018). If origin, breeding treatment and/or sex were involved in significant interactions, we calculated post-hoc contrasts on the estimated marginal means of their levels within levels of other factors involved in the respective interaction (R-package: emmeans v1.5.1, Lenth 2020). Multivariate statistical analyses of the full VOC-dataset are summarised in S9.

## 3. Results

### 3.1 Floral attractiveness

Spatial flower attractiveness of *S. latifolia* varied pronouncedly between plants of different breeding treatments, sexes, and population origins (Table 2). Inflorescences of inbreds were shorter than those of outbreds (p < 0.001, χ^2^_(1DF)_ = 37.31, Fig. 1a). Flower number (Fig. 1b) was higher in plants from North America than Europe (p = 0.005, χ^2^_(1DF)_ = 8.01) and additionally depended on the interaction breeding treatment × sex (p = 0.003, χ^2^_(1DF)_ = 8.99). Inbred plants generally produced fewer flowers than outbreds, and this effect was more severe in females (p_post_ < 0.001) than males (p_post_ = 0.011). The number of flowers produced was higher in male than female plants in both inbreds (p_post_ < 0.001) and outbreds (p_post_ < 0.001). The area of petal limbs (Fig. 1c) was smaller in female than male plants (p < 0.001, χ^2^_(1DF)_ = 51.35) and reduced by inbreeding (p = 0.002, χ^2^_(1DF)_ = 9.25). The expansion of the corolla depended on the interaction breeding treatment × sex (p = 0.004, χ^2^_(1DF)_ = 8.17). Inbreeding reduced corolla expansion in female (p_post_ < 0.001) but not in male plants, and differences between sexes in corolla expansion were consequently apparent in inbreds (p_post_ < 0.001) but not in outbreds (Fig. 1d). Corolla expansion additionally depended on the interaction sex × origin (p = 0.009, χ^2^_(1DF)_ = 6.86). Flowers expanded larger in male than female plants in populations originating from North America only (p_post_ < 0.001).

**Table 2.** Overview and results of statistical analyses with (generalised) linear mixed effects models. The table summarises the model types and error distributions used for each of the responses (printed in subscript), the parameter estimates on the link function scale with significance levels assessed based on Wald χ^2^ tests for all fixed effects (***: p < 0.001, **: p < 0.01, and *p < 0.05 printed in bold; Ns.: p > 0.05), and random effect variances (printed in italic). For zero-inflated responses, estimates from the conditional model parts appear in the first line and estimates from zero-inflation model parts in the second line.

|  | Intercept | Btmt<br>[outbred -<br>inbred] | Sex<br>[female -<br>male] | Origin<br>[Europe<br>- US] | Btmt<br>×<br>sex | Btmt<br>×<br>origin | Sex<br>×<br>origin | Btmt<br>×<br>sex<br>×<br>origin | Latitude | Plant<br>age | Obs. time | Pop | Pop<br>:<br>mother | Pop<br>:<br>father | Plot | Obs.<br>trial |
| --- | --- | --- | --- | --- | --- | --- | --- | --- | --- | --- | --- | --- | --- | --- | --- | --- |
| **Spatial flower traits** |  |  |  |  |  |  |  |  |  |  |  |  |  |  |  |  |
| Plant height<sub>LMM(G)</sub> | 81.03 | **3.56\*\*\*** | -0.40<sup>Ns</sup> | 1.84<sup>Ns</sup> | 0.67<sup>Ns</sup> | 0.11<sup>Ns</sup> | 1.07<sup>Ns</sup> | -0.15<sup>Ns</sup> | 1.00<sup>Ns</sup> | Nt. | Nt. | 5.42 | 16.76 | 0.00 | Nt. | Nt. |
| No. flowers<sub>GLMM(NBQ)</sub> | 2.61 | **0.14\*\*\*** | **-0.70\*\*\*** | **-0.25\*\*** | **0.08\*\*** | -0.04<sup>Ns</sup> | 0.01<sup>Ns</sup> | 0.01<sup>Ns</sup> | 0.12<sup>Ns</sup> | Nt. | Nt. | 0.03 | 0.02 | 0.02 | Nt. | Nt. |
| Petal limb area<sub>LMM(G)</sub> | 2.76 | **0.16\*\*** | **-0.38\*\*\*** | 0.07<sup>Ns</sup> | 0.06<sup>Ns</sup> | 0.04<sup>Ns</sup> | 0.10<sup>Ns</sup> | 0.03<sup>Ns</sup> | 0.19<sup>Ns</sup> | Nt. | Nt. | 0.15 | 0.08 | 0.00 | Nt. | Nt. |
| Corolla expansion<sub>LMM(G)</sub> | 5.21 | **0.21\*\*** | **-0.44\*\*\*** | 0.33<sup>Ns</sup> | 0.23<sup>Ns</sup> | 0.02<sup>Ns</sup> | **0.21\*\*** | 0.11<sup>Ns</sup> | 0.06<sup>Ns</sup> | Nt. | Nt. | 0.29 | 0.30 | 0.00 | Nt. | Nt. |
| **Flower scent composition** |  |  |  |  |  |  |  |  |  |  |  |  |  |  |  |  |
| Shannon index VOC<sub>LMM(G)</sub> | 1.86 | 0.03<sup>Ns</sup> | **-0.13\*** | -0.06<sup>Ns</sup> | 0.01<sup>Ns</sup> | 0.04<sup>Ns</sup> | 0.02<sup>Ns</sup> | 0.01<sup>Ns</sup> | 0.03<sup>Ns</sup> | **-0.02\*** | Nt. | 0.01 | 0.00 | 0.00 | Nt. | Nt. |
| Lilac aldehyde A<sub>ZI-GLMM(NBQ)</sub> | 15.11 | 0.02<sup>Ns</sup> | -0.06<sup>Ns</sup> | 0.02<sup>Ns</sup> | 0.01<sup>Ns</sup> | -0.08<sup>Ns</sup> | -0.04<sup>Ns</sup> | **-0.16\*** | 0.01<sup>Ns</sup> | -0.08 | Nt. | 0.01 | 0.00 | 0.00 | Nt. | Nt. |
|  | -1.93*** | Nt. | Nt. | Nt. | Nt. | Nt. | Nt. | Nt. | Nt. | Nt. | Nt. | Nt. | Nt. | Nt. | Nt. | Nt. |
| Lilac aldehyde B/C<sub>ZI-GLMM(NBL)</sub> | 15.78 | -0.08<sup>Ns</sup> | -0.09<sup>Ns</sup> | 0.12<sup>Ns</sup> | 0.06<sup>Ns</sup> | -0.10<sup>Ns</sup> | -0.01<sup>Ns</sup> | -0.03<sup>Ns</sup> | -0.09<sup>Ns</sup> | -0.03<sup>Ns</sup> | Nt. | 0.01 | 0.00 | 0.03 | Nt. | Nt. |
|  | -1.93*** | Nt. | Nt. | Nt. | Nt. | Nt. | Nt. | Nt. | Nt. | Nt. | Nt. | Nt. | Nt. | Nt. | Nt. | Nt. |
| Lilac aldehyde D<sub>ZI-GLMM(NBQ)</sub> | 13.70 | -0.05<sup>Ns</sup> | 0.14<sup>Ns</sup> | -0.07<sup>Ns</sup> | -0.09<sup>Ns</sup> | -0.16<sup>Ns</sup> | -0.06<sup>Ns</sup> | -0.08<sup>Ns</sup> | 0.08<sup>Ns</sup> | -0.17<sup>Ns</sup> | Nt. | 0.00 | 2.75 | 0.47 | Nt. | Nt. |
|  | 0.02<sup>Ns</sup> | Nt. | Nt. | Nt. | Nt. | Nt. | Nt. | Nt. | Nt. | Nt. | Nt. | Nt. | Nt. | Nt. | Nt. | Nt. |
| **Flower colour** |  |  |  |  |  |  |  |  |  |  |  |  |  |  |  |  |
| Reflectance UV<sub>LMM(G)</sub> | 2.04 | -0.05<sup>Ns</sup> | **-0.27\*\*\*** | 0.05<sup>Ns</sup> | 0.04<sup>Ns</sup> | 0.08<sup>Ns</sup> | -0.04<sup>Ns</sup> | 0.02<sup>Ns</sup> | 0.01<sup>Ns</sup> | Nt. | Nt. | 0.01 | 0.03 | 0.01 | Nt. | Nt. |
| Reflectance blue<sub>LMM(G)</sub> | 26.72 | -0.02<sup>Ns</sup> | **-0.95\*\*\*** | 0.10<sup>Ns</sup> | 0.24<sup>Ns</sup> | 0.05<sup>Ns</sup> | 0.21<sup>Ns</sup> | 0.14<sup>Ns</sup> | 0.47<sup>Ns</sup> | Nt. | Nt. | 0.47 | 0.85 | 0.00 | Nt. | Nt. |
| Reflectance green<sub>LMM(G)</sub> | 39.87 | -0.10<sup>Ns</sup> | -0.19<sup>Ns</sup> | 0.40<sup>Ns</sup> | 0.26<sup>Ns</sup> | 0.17<sup>Ns</sup> | 0.12<sup>Ns</sup> | 0.31<sup>Ns</sup> | 0.05<sup>Ns</sup> | Nt. | Nt. | 1.12 | 0.37 | 0.07 | Nt. | Nt. |
| **Floral rewards** |  |  |  |  |  |  |  |  |  |  |  |  |  |  |  |  |
| Excreted sugar<sub>LMM(G)</sub> | 5.70 | 0.03<sup>Ns</sup> | **-0.18\*\*\*** | -0.08<sup>Ns</sup> | 0.04<sup>Ns</sup> | -0.02<sup>Ns</sup> | -0.02<sup>Ns</sup> | 0.04<sup>Ns</sup> | 0.04<sup>Ns</sup> | Nt. | Nt. | 0.07 | 0.00 | 0.00 | Nt. | Nt. |
| **Pollinator visitation** |  |  |  |  |  |  |  |  |  |  |  |  |  |  |  |  |
| No. plant visits<sub>ZI-GLMM(P)</sub> | 0.28 | 0.09<sup>Ns</sup> | **-0.67\*\*\*** | 0.09<sup>Ns</sup> | 0.22<sup>Ns</sup> | -0.20<sup>Ns</sup> | -0.16<sup>Ns</sup> | **-0.29\*** | Nt. | Nt. | **-0.11\*** | Nt. | Nt. | Nt. | 0.27 | 0.15 |
|  | -0.48<sup>Ns</sup> | 0.05<sup>Ns</sup> | 0.29<sup>Ns</sup> | -0.21<sup>Ns</sup> | -0.11<sup>Ns</sup> | -0.11<sup>Ns</sup> | 0.10<sup>Ns</sup> | **-0.46\*** | Nt. | Nt. | 0.10<sup>Ns</sup> | Nt. | Nt. | Nt. | 0.14 | 0.44 |
| No. flowers visited<sub>ZI-GLMM(P)</sub> | 0.79 | 0.04<sup>Ns</sup> | **-0.56\*\*\*** | -0.03<sup>Ns</sup> | **0.34\*** | -0.23<sup>Ns</sup> | -0.22<sup>Ns</sup> | **-0.46\*\*** | Nt. | Nt. | **0.19\*\*\*** | Nt. | Nt. | Nt. | 0.68 | 0.23 |
|  | -0.09<sup>Ns</sup> | -0.04<sup>Ns</sup> | **0.47\*\*** | -0.23<sup>Ns</sup> | -0.19<sup>Ns</sup> | -0.02<sup>Ns</sup> | 0.17<sup>Ns</sup> | **-0.31\*** | Nt. | Nt. | **0.47\*\*** | Nt. | Nt. | Nt. | 0.16 | 0.47 |

Abbreviations. Btmt: breeding treatment, LMM: linear mixed effects model, (NBQ): negative binomial distribution with quadratic parametrization and log-link, (NBL): negative binomial distribution with linear parametrization and log-link, No.: number, Nt.: not tested, GLMM: generalised linear mixed effects model, Obs: Observation, (P): Poisson distribution with log-link, Pop: population, ZI-GLMM: zero-inflation generalised mixed effects model, (G): Gaussian distribution with identity link

**Fig. 1:**
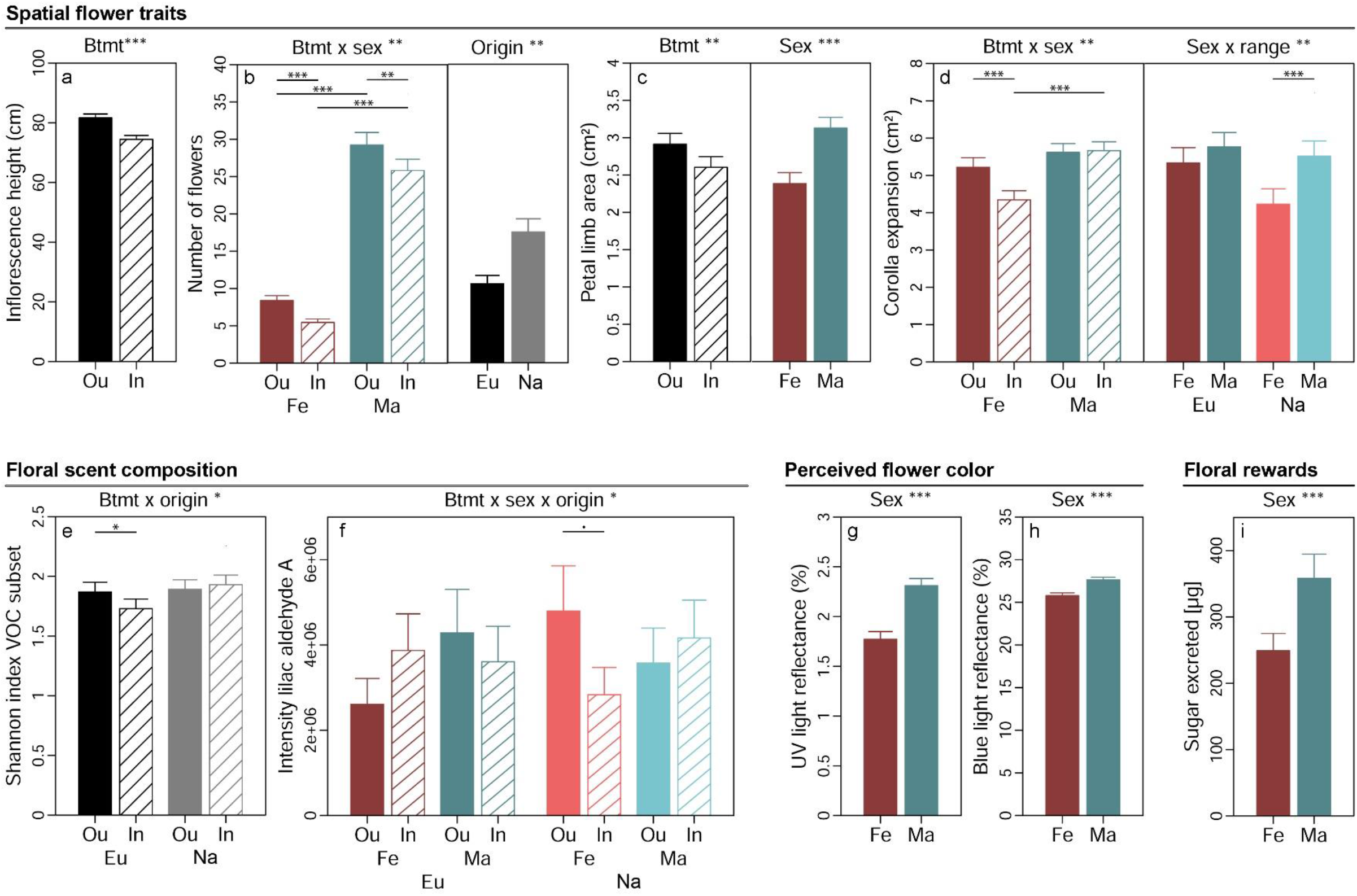
Effects of breeding treatment, sex and origin on spatial flower attractiveness traits (a-d), flower scent composition (e-f), flower colour as perceived by crepuscular moths (g-h) and floral rewards in *Silene latifolia*. Graphs show estimated marginal means and standard errors for outbred (Ou, filled bars) and inbred (In, open bars), female (Fe, red bars) and male (Ma, blue bars) plants from Europe (Eu, dark coloured bars) and the North America (Na, bright coloured bars). Estimates were extracted from (generalised) linear mixed effects models for significant interaction effects and main effects of factors not involved in an interaction (significance levels based on Wald χ^2^-tests denoted at top of plot). Interaction effect plots additionally indicate significant differences among breeding treatments, sexes or origins within levels of other factors involved in the respective interaction (estimated based on post-hoc comparisons, denoted within plots). Exact sample sizes for all traits are listed in Table 1. Significance levels: ***: p < 0.001, **: p < 0.01, *p < 0.05, • p < 0.06.

Breeding treatment, sex and population origin affected on the composition of floral VOC in *S. latifolia* interactively (Table 2). The Shannon diversity of VOC known to elicit antennal responses in *H. bicruris* depended on the interaction breeding treatment × origin (p = 0.016, χ^2^_(1DF)_ = 5.83, Fig. 1e). Inbreeding slightly reduced Shannon diversity of these VOC in European plants (p_post_ = 0.013) but had no effect on plants from North America. The intensity of lilac aldehyde A depended on the interaction breeding treatment × sex × origin in the conditional model (p = 0.025, χ^2^_(1DF)_ = 5.03, Fig. 1f). Post-hoc comparisons yielded a marginally significant lower intensity of the compound in inbred than outbred females in plants from North America (p_post_ = 0.056). Similar non-significant trends were observed for the other lilac aldehyde isomers (Table S5). Multivariate statistical analyses of all 70 VOC revealed no clear separation of floral headspace VOC patterns for any of the treatments (S8).

The proportion of flower colour detectable for crepuscular moths and the sugar excreted as reward with nectar were independent of breeding treatment and population origin but exhibited differences between plants of different sex (Table 2). Male flowers reflected more light in the spectrum detectable by the UV receptor (350 nm) (p < 0.001, χ^2^_(1DF)_ = 41.92, Fig. 1g) and the blue receptor (440 nm) (p < 0.001, χ^2^_(1DF)_ = 39.59, Fig. 1h) than flowers of females. Likewise, the amount of sugar excreted with nectar was higher in male than female plants (p < 0.001, χ^2^_(1DF)_ = 14.16, Fig. 1i).

### 3.2 Pollinator visitation rates

The number of pollinator visits per plant by moths was shaped by the interaction breeding treatment × sex × origin in the conditional model (p = 0.016, χ^2^_(1DF)_ = 5.84, Table 2, Fig. 2a). Post-hoc comparisons yielded that plant visits were reduced by inbreeding in female plants from North America (p_post_ = 0.007); higher in male than female plants in European outbreds (p_post_ < 0.001), European inbreds (p_post_ = 0.001), and North American inbreds (p_post_ < 0.001); and higher in plants from North America than Europe in outbred females (p_post_ = 0.014). The number of flowers approached per plant visit was likewise shaped by the interaction breeding treatment × sex × origin in the conditional model (p = 0.001, χ^2^_(1DF)_ = 10.61, Fig. 2b, Table 2). Post-hoc comparisons yielded that flower visits were lower for inbred females (p_post_ = 0.001) but higher for inbred males (p_post_ = 0.031) in populations from North America; higher in males than females for European outbreds (p_post_ = 0.003), European inbreds (p_post_ = 0.027) and North American inbreds (p_post_ = 0.002) but higher in females than males in outbreds from North America (p_post_ < 0.001); and higher in North American than European outbred females (p_post_ = 0.001) (Fig 2b). Both, the number of plant and flower visits depended on the interaction breeding treatment × sex × origin in the zero-inflation part of the model as well (Table 2, S9). The direction and magnitude of these effects did not contrast with the conditional models.

**Fig. 2:**
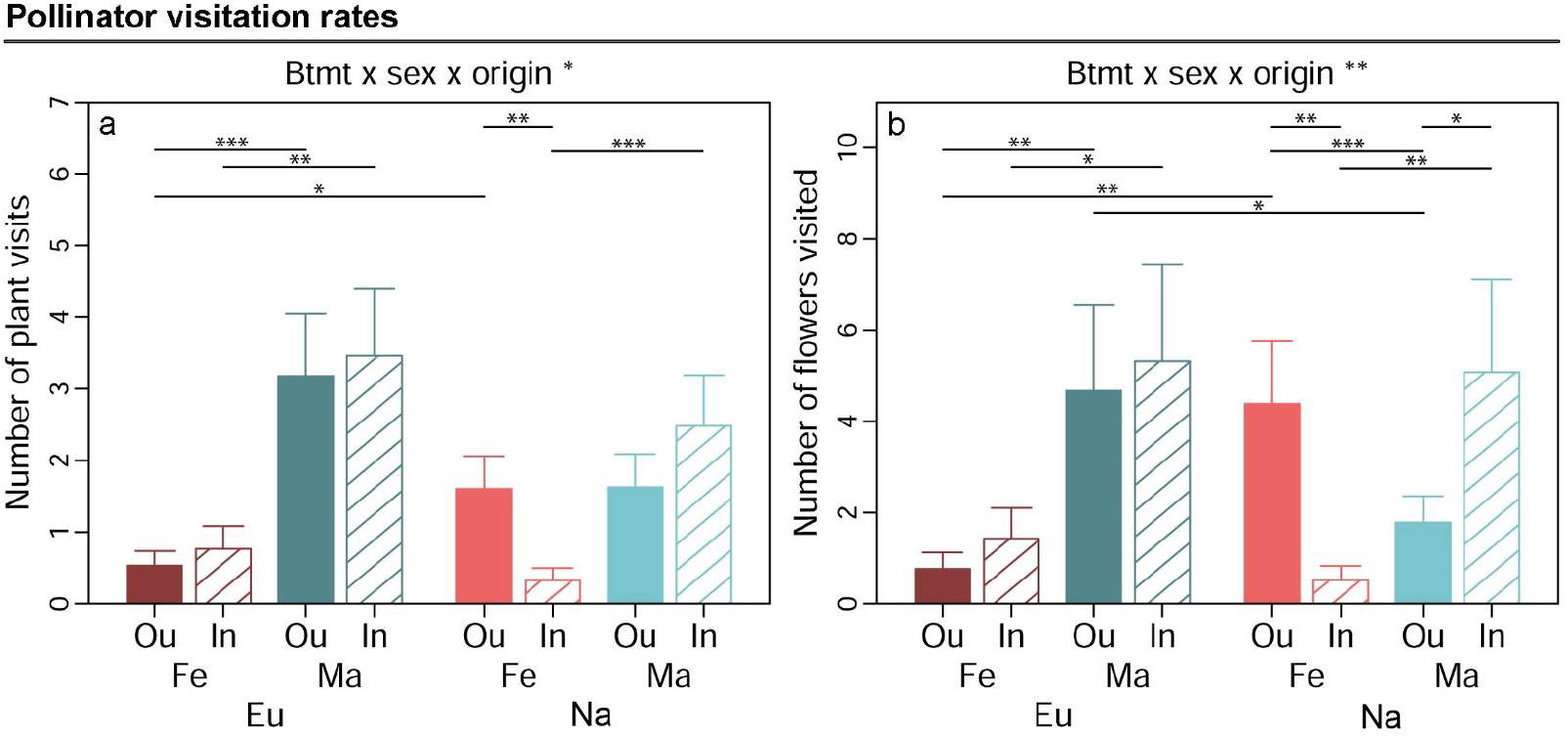
Effects of breeding treatment, sex and origin on pollinator visitation rates in *Silene latifolia*. Graphs show estimated marginal means and standard errors for outbred (Ou, filled bars) and inbred (In, open bars), female (Fe, red bars) and male (Ma, blue bars) plants from Europe (Eu, dark coloured bars) and North America (Us, bright coloured bars). Estimates were extracted for significant interaction effects from the conditional part of generalised linear mixed effects models (significance levels based on Wald χ^2^-tests denoted at top of plot). Plots additionally indicate significant differences between breeding treatments, sexes or origins within levels of other factors involved in the respective interaction (estimated based on post-hoc comparisons, denoted within plots). Exact sample sizes for all traits are listed in Table 1. Significance levels: ***: p < 0.001, **: p < 0.01, and *p < 0.05.

## 4. Discussion

Using an integrated methodological approach, we found that i) inbreeding diminishes several flower attractiveness traits in *S. latifolia*. The magnitude of these effects depended partially on ii) plant sex, which demonstrates that the intrinsic biological differences between males and females shape the consequences of inbreeding in dioecious plant species as they are filtered through the selective environment. Inbreeding effects also depended on iii) origin in a way indicating that divergent evolutionary histories have shaped the underlying genetic architecture. Finally, our study showed that iv) the effects of inbreeding, sex and origin on pollinator visitation rates specifically mirrored variation in floral scent, which highlights the large relative importance of this attractiveness trait in shaping the behaviour of crepuscular moths.

### 4.1 Inbreeding reduces floral attractiveness

In partial accordance with our first hypothesis, inbreeding reduced several, but not all components of floral attractiveness in *S. latifolia*. Spatial attractiveness declined most strongly with inbreeding in males and females from both origins (Fig. 1a-c). These results are in line with previous studies on hermaphroditic, self-compatible species (Ivey and Carr, 2005; Glaettli and Goudet, 2006) and support that the complex genetic architecture underlying such traits (Feng et al., 2019) gives rise to dominance and over-dominance effects at multiple loci. The diversity and quantity of floral VOC was reduced in a sex and origin-specific manner in inbred relative to outbred *S. latifolia* (Fig. 1e-g). So far, lower emissions of floral volatiles in inbreds have been reported for only few plants species pollinated by diurnal generalists (Ferrari et al., 2006; Haber et al., 2019). Our study demonstrates such effects for plants pollinated by specialist moths that use scent as a major cue for plant location (Riffell and Alarcón, 2013). Dominance and over-dominance may either have directly interfered with genes involved in VOC-syntheses and their regulation in *S. latifolia* or unfolded their effects by disrupting physiological homoeostasis and thereby inducing intrinsic stress that came at the cost of scent production (Kristensen et al., 2010; Fox and Reed, 2011).

In contrast to spatial structure and scent, flower colour and the total amount of sugar excreted with nectar exhibited no differences among inbreds and outbreds in *S. latifolia* (Table 2). Flower colour is an attractiveness trait that has, to our knowledge, not yet been studied in the context of inbreeding, despite its crucial role in flower identification and localization (Garcia et al., 2019). We demonstrate that the flower colour perceived by moths is not altered by inbreeding and seems to be a conserved trait in *S. latifolia*. Other species with high intraspecific variation in flower colour may be ideal models to further examine the relationship with inbreeding in the future by combining visual modelling with choice experiments (Kelber et al., 2003). The independence of sugar excretion from inbreeding in *S. latifolia* indicates that the strong reduction of flower number in inbreds allows compensating the quality of floral rewards *via* a resource allocation trade-off.

Overall, the observed inbreeding effects on attractiveness were partially small and variable in their magnitude as compared to previous investigations. However, our findings highlight that even weak degrees of biparental inbreeding (i.e., one generation sib-mating) can result in a comprehensive reduction of spatial and olfactory flower attractiveness that is detectable against the background of natural variation among multiple plant populations from a broad geographic region. Most importantly, variation in inbreeding effects was consistent in its dependency on plant sex, which gives insight into the role of intrinsic biological differences between males and females in the expression of inbreeding depression.

### 4.2 The cost of inbreeding for floral attractiveness is higher in females than males

Males outperformed females in all attractiveness traits, except scent production (Fig. 1). As such, our study confirmed previously observed sexual dimorphisms in *S. latifolia* (nectar: Gehring et al. 2004; flower number: Delph et al. 2010) but also yielded contradicting results. As opposed to Delph et al. (2010), we observed larger instead of smaller flowers in males. This may base on the use of a size estimate that accounts for variation in flower shape or the comparably large geographic range and higher number of populations covered by our study. Moreover, we discovered a novel sexually dimorphic trait in the colour appearance of *S. latifolia* to crepuscular moths in the UV and blue light spectrum (Fig. 1g-h). Given that moths use blue light as a major cue to start feeding on nectar (Cutler et al., 1995), the lower light reflectance observed for females is another trait rendering them less attractive than males.

The evolution of lower female attractiveness is driven by sex-specific resource allocation, i.e., high costs of ovary and seeds restrict allocation to floral attractiveness in females, as well as sexual selection, i.e., competition for siring success among males selects for increased attractiveness (Moore & Pannell 2011; Barrett & Hough 2013). Both processes may also explain the larger magnitude of inbreeding effects in female plants of *S. latifolia*, which we observed in accordance with our second hypothesis (Fig. 1b, d). High reproductive expenditure in females may increase the frequency and intensity of resource depletion stress (e.g., drought) under field conditions (Obeso, 2002; Li et al., 2004; Zhang et al., 2010). Consequently, females may suffer disproportionally from inbreeding when dominance and over-dominance affect loci mediating resistance to such stress (Fox and Reed, 2011). The assumption that sex-specific viability selection plays a role in the expression of inbreeding depression seems likely for *S. latifolia*. Previous research elaborated that females of this species experience resource depletion stress more often than males in the late growing season during fruit maturation (Gehring and Monson, 1994) and that inbreeding reduces resistance to environmental stress (Schrieber et al., 2018, 2019). Another non-exclusive explanation for the different inbreeding effects on female versus male *S. latifolia* may be found in sexual selection. Competition for increased siring success among males could rapidly purge deleterious recessive mutations in genes directly linked to attractiveness. Finally, a proportion of sex-specific inbreeding effects may be attributed to differential gene expression. *Silene latifolia* harbours numerous genes with alleles that affect male and female fitness in opposite directions. These sexually antagonistic genes are partly subsumed in non-recombining regions of gonosomes (Scotti and Delph, 2006). Those located at the X-chromosome are always effectively dominant in males (XY) but may be recessive and therefore contribute to inbreeding depression in females (XX). The remaining fraction of sexually antagonistic genes is located at autosomal regions but exhibits sex-specific expression as controlled by the gonosomes (Scotti and Delph, 2006). These genes may exhibit systematic differences in the abundance and effect magnitudes of deleterious recessive alleles between males and females, thus contributing to sex-specific inbreeding effects.

Not only flower attractiveness but also plant viability may exhibit sex-specific inbreeding depression in dioecious species. This could result in deviations from optimal sex ratio and, consequently, reductions of effective population sizes that accelerate local extinctions under global change (Hultine et al., 2016; Rosche et al., 2018). Future studies should aim at disentangling the relative contribution of sex-specific selection and gene expression to differences in the magnitude of inbreeding depression between males and females and at predicting their feedback on sex ratios to predict and handle these specific threats.

### 4.3 Evolutionary history shapes the genetic architecture underlying inbreeding effects on floral scent

Plants exhibited merely one general difference in attractiveness among geographic origins (Fig. 1b). Indeed, we had not expected broad differences in floral attractiveness among European and North American *S. latifolia* plants (i.e., significant main effects of origin). A sufficient overlap in the composition of pollinator communities (Castillo et al., 2014) and appropriate pre-adaptations in attractiveness were probably essential for *S. latifolia* as an obligate outcrossing plant species to successfully colonise North America. Even the higher flower numbers in North American *S. latifolia* plants (Fig. 1b) do not necessarily result from pollinator-mediated selection but rather from changes in the selective regimes for numerous abiotic factors (Keller et al., 2009).

However, we expected that North American populations purged genetic load linked to floral attractiveness during the colonization process (i.e., interaction breeding treatment × origin). In contrast to hypothesis three, the magnitude of inbreeding effects was not consistently higher in European than North American populations. Instead, it was independent of origin for most attractiveness components, except flower scent, and either higher or lower in European plants for different scent traits (Fig. 1f-g). These findings provide no support for recent purging-events in North American populations. They rather add to evidence that the magnitude of inbreeding effects is highly specific for the traits as well as the populations or population groups under investigation (e.g., Escobar et al., 2008; Angeloni et al., 2011). This specifity roots in the composition of gene-loci affected by dominance and overdominance and is determined by the complex interplay of demographic population histories (i.e., size retractions and expansions, genetic drift, isolation, gene flow) and the selective environment (Charlesworth and Willis, 2009). As such, the precise mechanisms underlying variation in inbreeding effects on different scent traits across population origins of *S. latifolia* can only be explored based on comprehensive genomic resources, which are currently not available.

### 4.4 Inbreeding effects on flower scent cause feedback on pollinator visitation rates

Pollinator visitation rates partially mirrored the above-discussed variation in flower attractiveness. They depended on the breeding treatment in a highly sex-and origin-specific manner: In North American populations, inbred females received significantly fewer plant and flower visits than outbreds, whereas flower visits were higher in inbred males (Fig. 2). Although these findings provide limited support for our fourth hypotheses, they yield interesting insight into the relative importance of single attractiveness traits for the behaviour of a lepidopteran specialist pollinator.

We conclude that the severe inbreeding effects on spatial attractiveness alone do not necessarily reduce moth visitation rates because these effects were independent of plant sex and origin. A trait that was negatively affected by inbreeding only in North American females, just like pollinator visitation rates, was the concentration of lilac aldehyde A (Fig. 1f). The other lilac aldehyde isomers exhibited similar but non-significant trends (Table S5). Indeed, electrophysiological experiments have shown that the antennae of *H. bicruris* can detect slight differences in lilac aldehyde concentrations at very low dosages (Dötterl et al. 2006). Moreover, the compounds elicit oriented flight and landing responses in the moth more than any other VOC in the scent of *S. latifolia* flowers. Consequently, low lilac aldehyde concentrations may have resulted in a low attraction of moths to North American inbred females from the distance or to an immediate switch to more attractive plants following first probing in our experiment. The non-significant trend for higher relative amounts of lilac aldehyde in inbred than outbred males from North America (Table S5) could also explain the corresponding variation observed in flower visitation rates. Although the effects of inbreeding, sex and origin on floral VOC in *S. latifolia* were either weak or not significant, they were more clearly mirrored in pollinator visitation rates than any other attractiveness trait. This may generally apply for specialised moth pollinators that use floral scent as a major cue for the identification and location of their target plants (Riffell and Alarcón, 2013) and thus evolve more elaborate olfactory than visual perception systems (Hansson and Stensmyr, 2011; van der Kooi et al., 2021). Even studies on bumble bees have shown that reduced spatial attractiveness and rewards cannot explain the lower visitation of inbred target plants (Carr et al., 2014), which indicates a high relative importance of flower scent (or alternatively flower colour) for diurnal generalist pollinators as well.

In summary, our research on *S. latifolia* supports that in addition to inbreeding disrupting interactions with herbivores by changing plant leaf chemistry (Schrieber et al., 2018) it affects plant interactions with pollinators by altering flower chemistry. These threats to antagonistic and symbiotic plant-insect interactions may mutually magnify in reducing plant individual fitness (Ivey and Carr, 2005) and altering the dynamics of natural plant populations under global change.

## Acknowledgements

This study was funded by the program for the promotion of young female scientists of the Faculty of Mathematics and Natural Sciences of Kiel University. We warmly thank Ms. Koopmann, who kindly provided her private property as field site for pollinator observations. Tim Diekötter provided germination chambers and greenhouse space and Wolfgang Bilger supported us with light standards for digital image acquisition. Susanne Petersen, Stephan Doose, David Eder, Carolin Böttcher, Jorun Jess, Sarai Guadalupe Quezada-Jimenez, Verena Zajonc, Ann-Cathrin Voss and Pia Music provided technical assistance.

## Provision of data

All datasets supporting this article will be deposited in Dryad upon acceptance of this article for publication.

## Supporting Information

**Figure S1:**
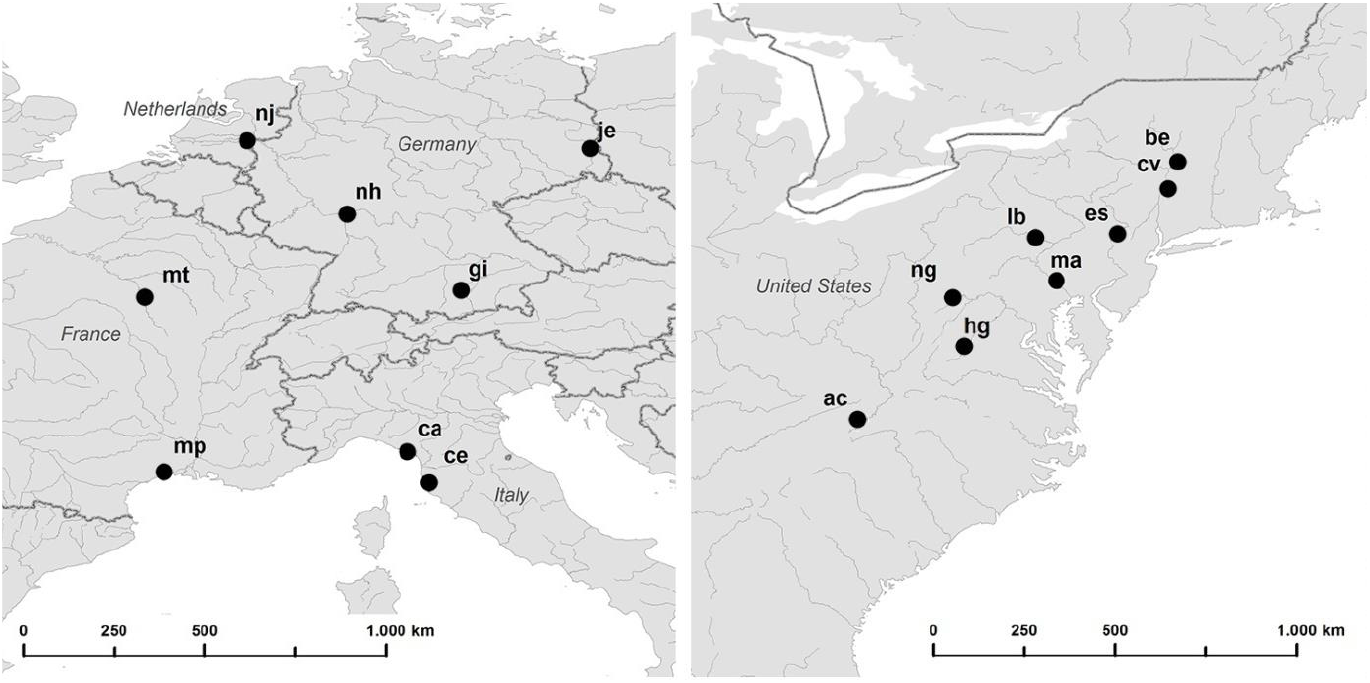
Map of the geographic locations of the sampled European (left) and North American (right) *Silene latifolia* populations.

**Table S1:**
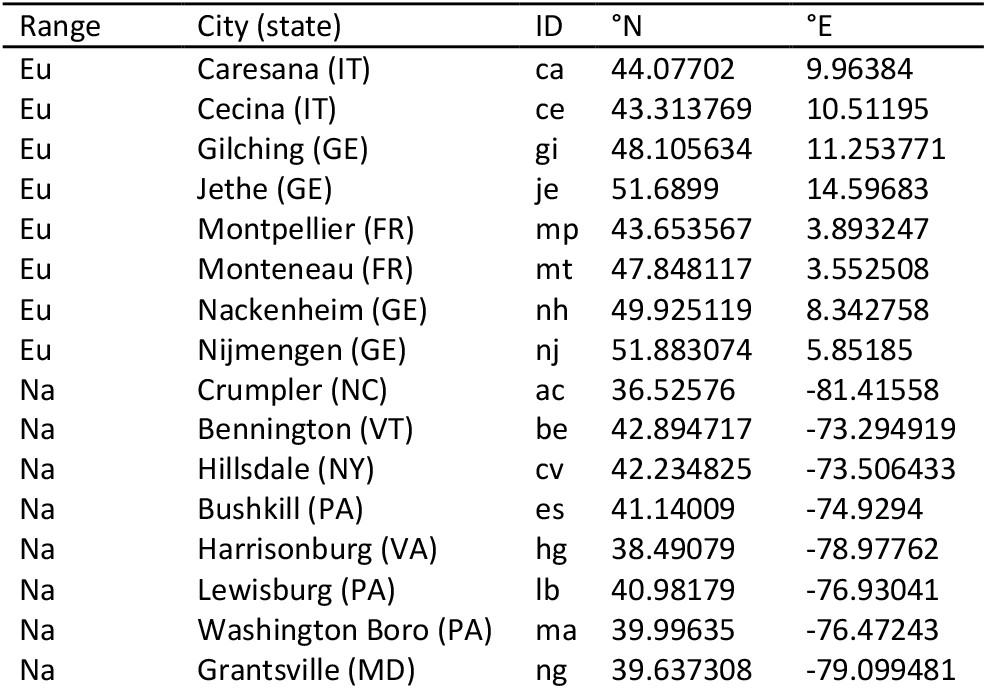
Overview of the geographic locations of *Silene latifolia* populations sampled in Europe (Eu) and the North America (Na).

**Figure S3:**
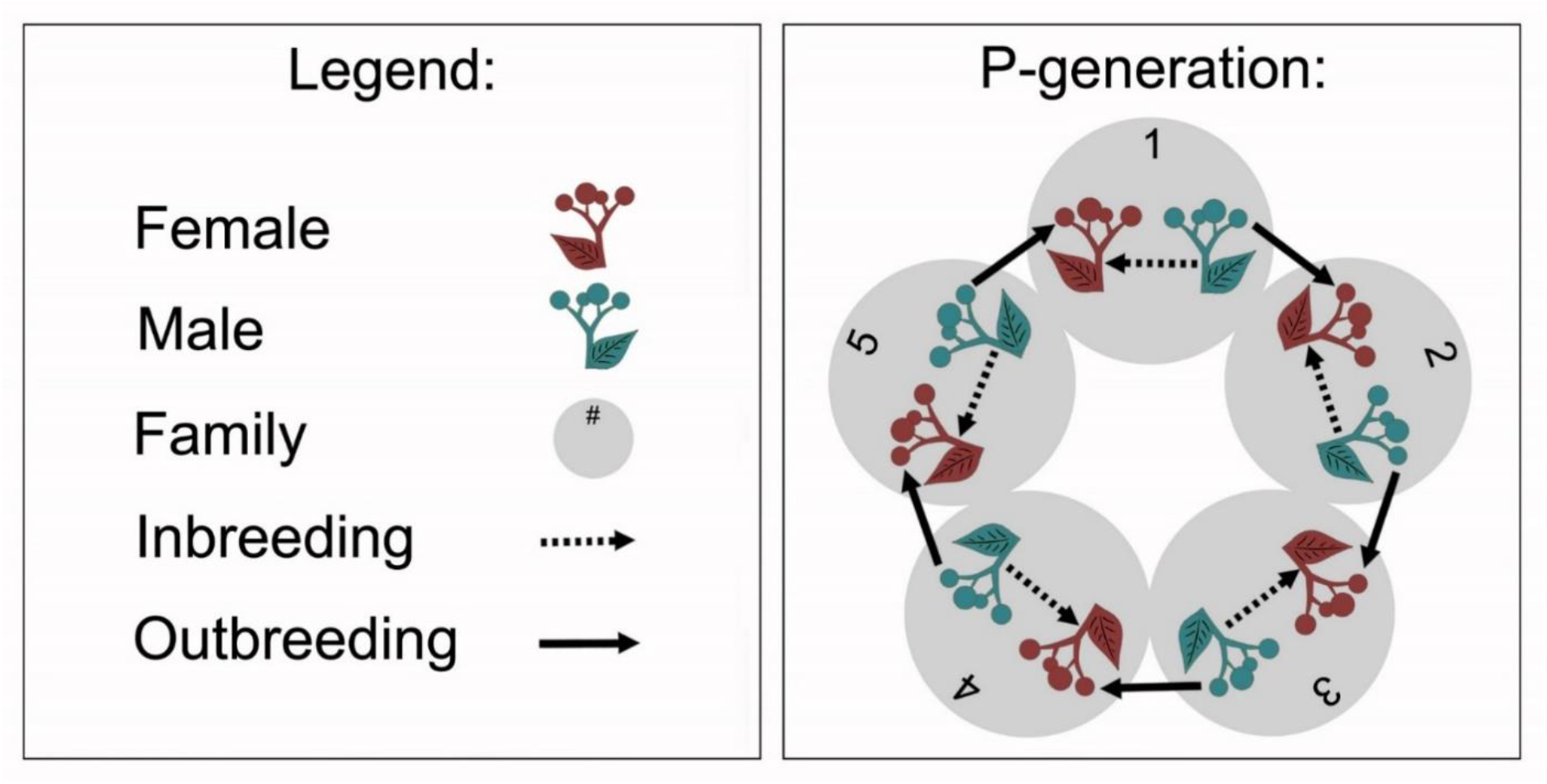
Overview of the experimental crossings within each of the 16 *Silene latifolia* populations. The crossings were performed with five families (numbered grey circles). Females (red plants, leaf pointing to left) were fertilised with pollen from males (blue plants, leaf pointing to right) from the same family for inbreeding (dashed arrows), and with pollen from males from a different family for outbreeding (solid arrows). Inbreeding and outbreeding were performed at distinct flowers of the same female individual.

**Figure S4:**
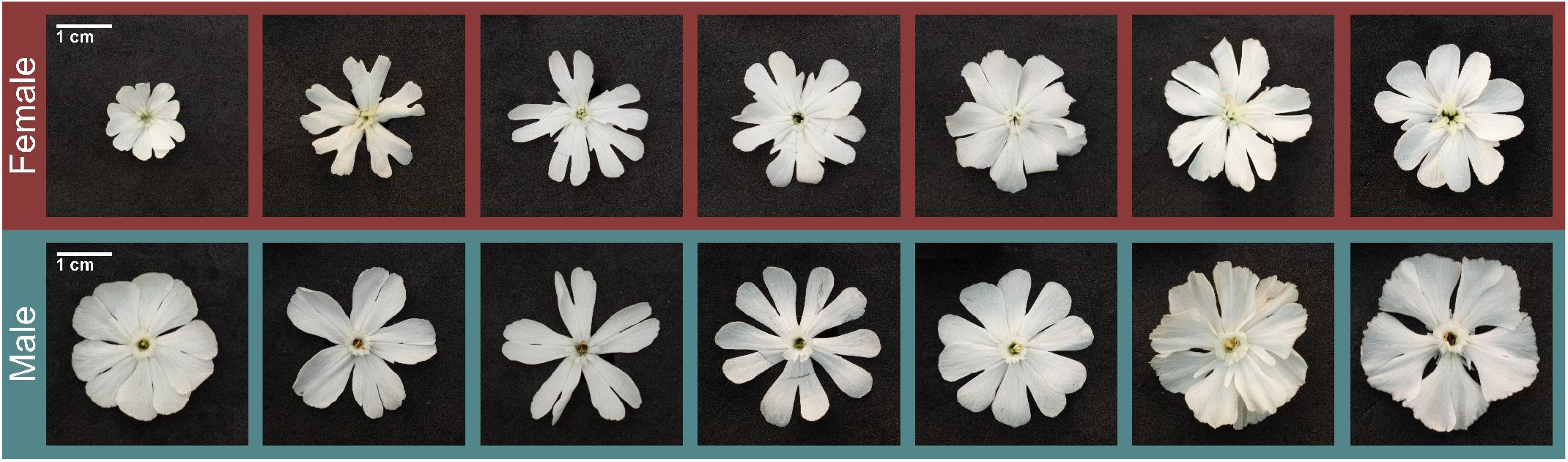
Variation in flower shape of *Silene latifolia* plants in our experiment. Photographs from female (upper row) and male (lower row) flowers with maximal deviation in flower shape were randomly chosen from the entire pool of experimental plants.

**Figure S5:**
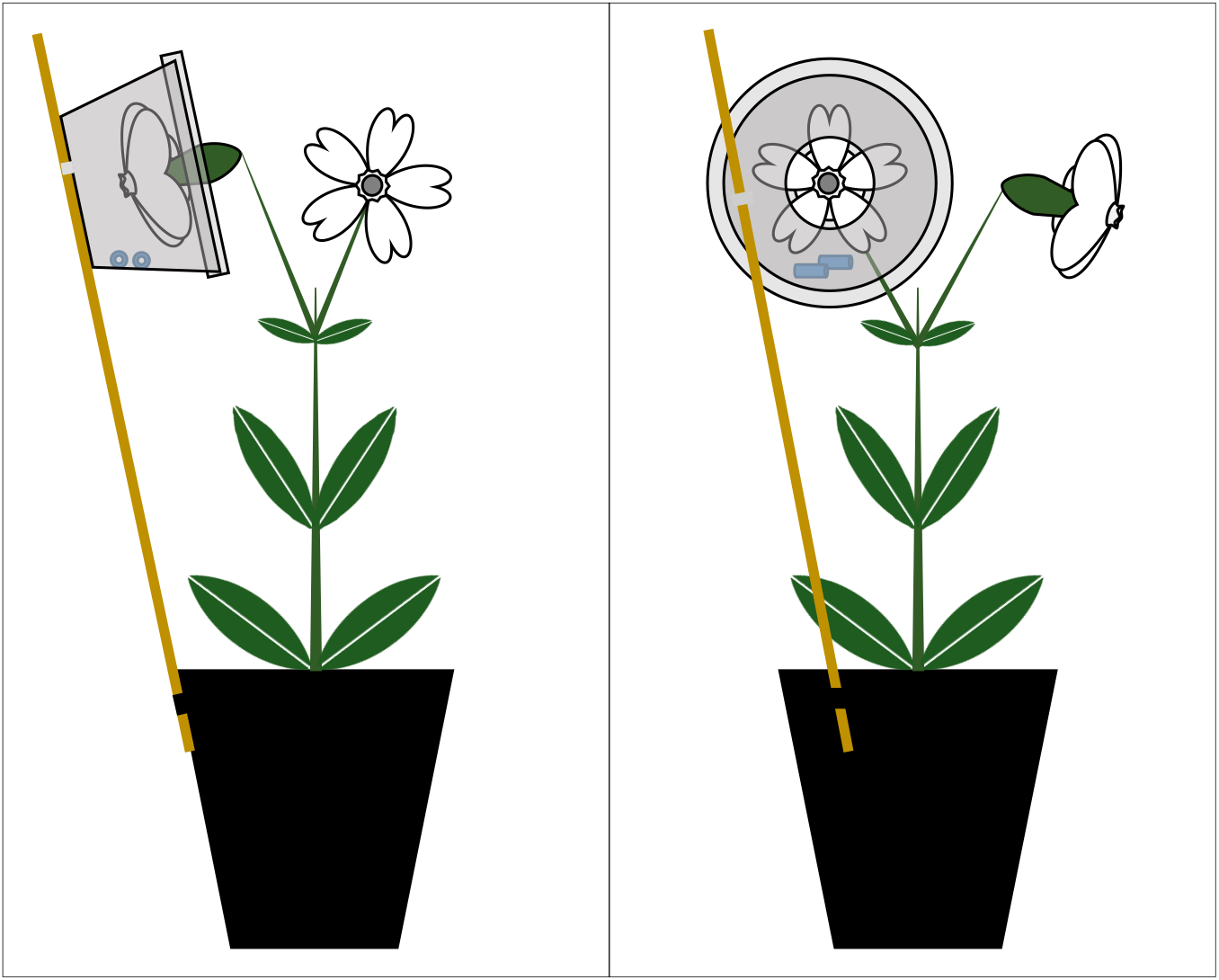
Experimental setup for the collection of headspace volatile organic compounds (VOC) from flowers of *Silene latifolia* (left: side view, right: front view). Flowers were inserted into VOC-collection units (consisting of 50 mL PE cups with lids, both with 15 mm holes), which were fixed *via* wooden sticks at the plant pot exterior. Two polydimethylsiloxane (silicone) tubes of standardised size were inserted into the collection units and absorbed VOC for a period of eight hours.

**Table S5:**
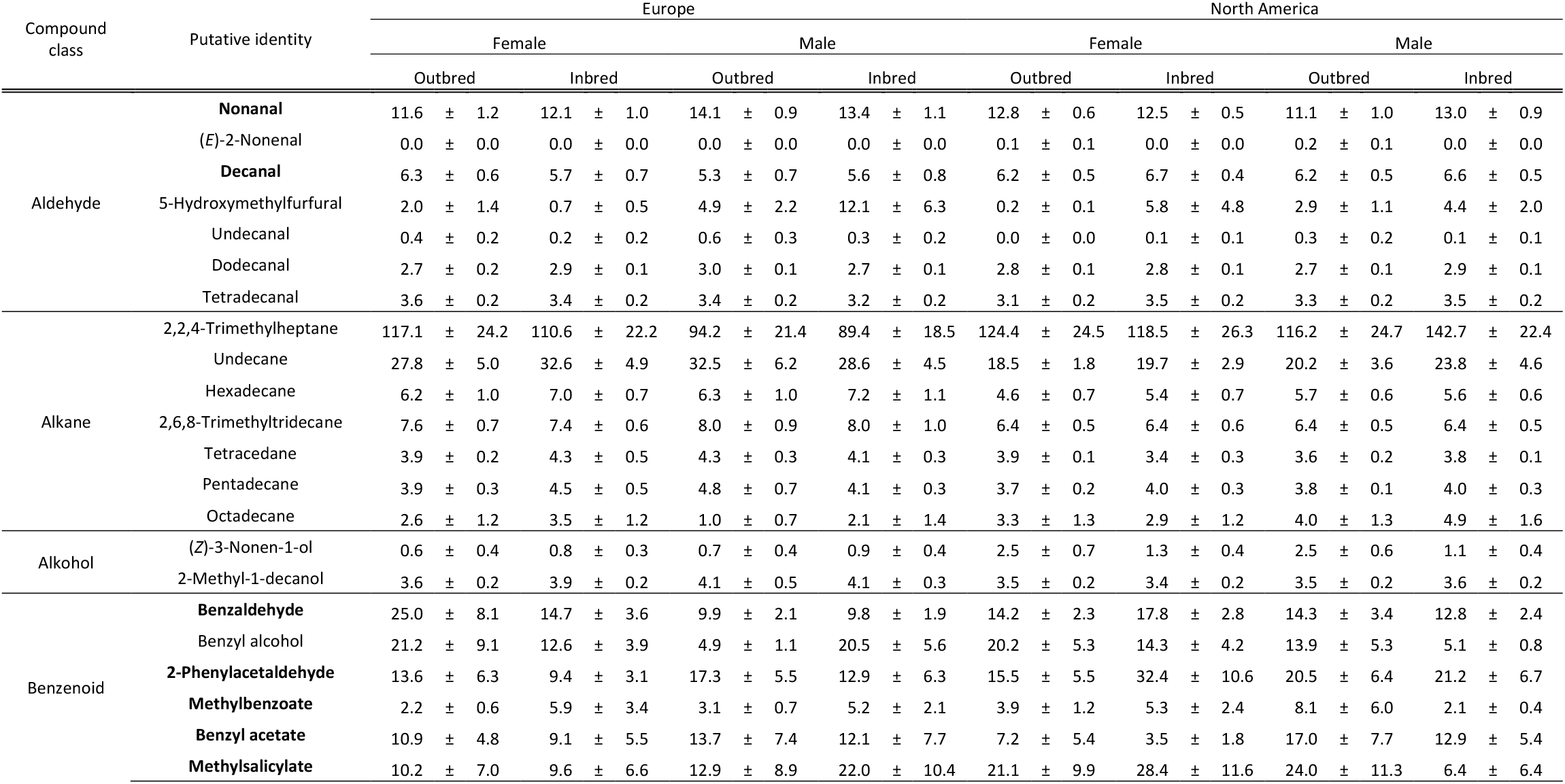
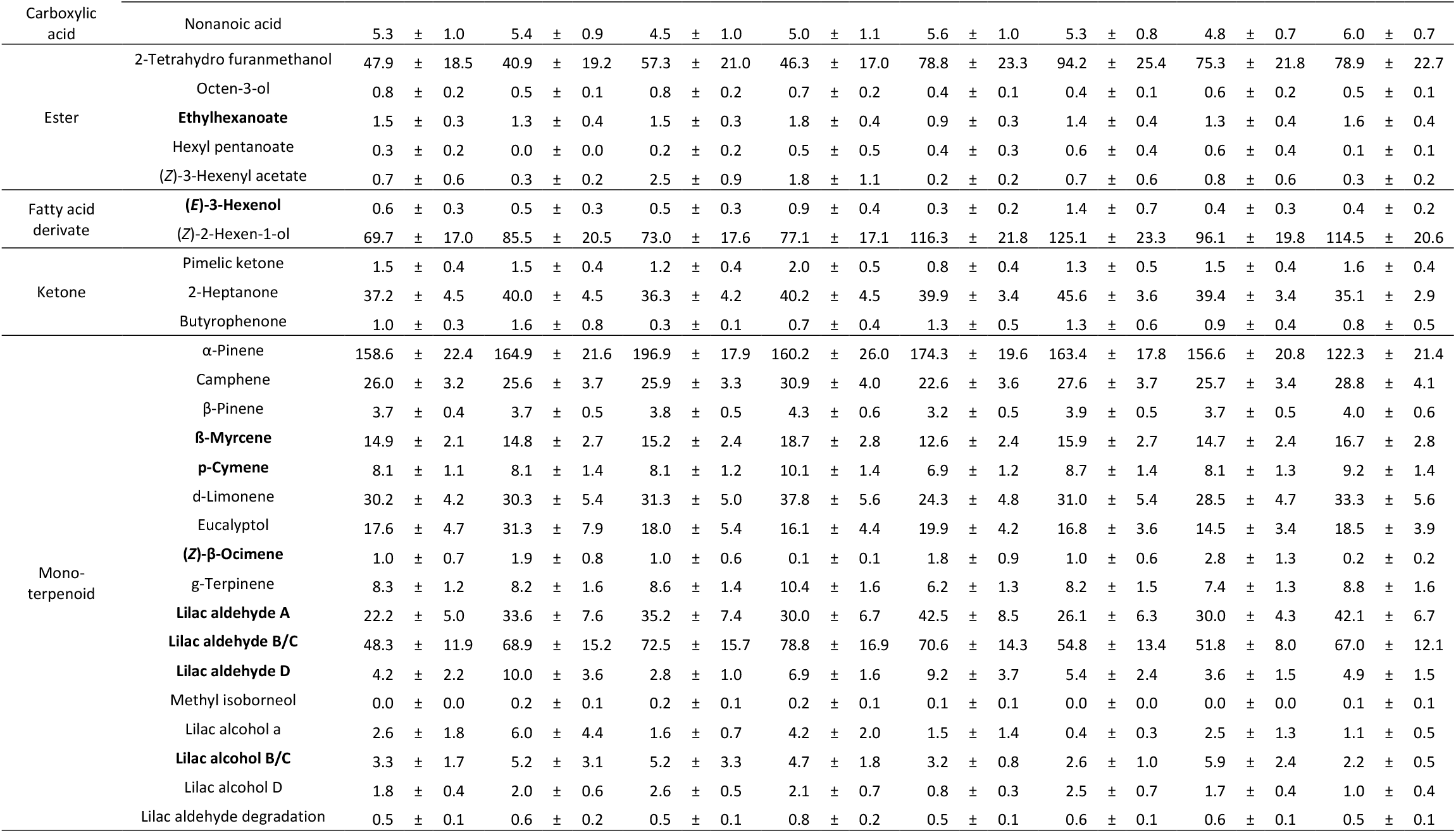
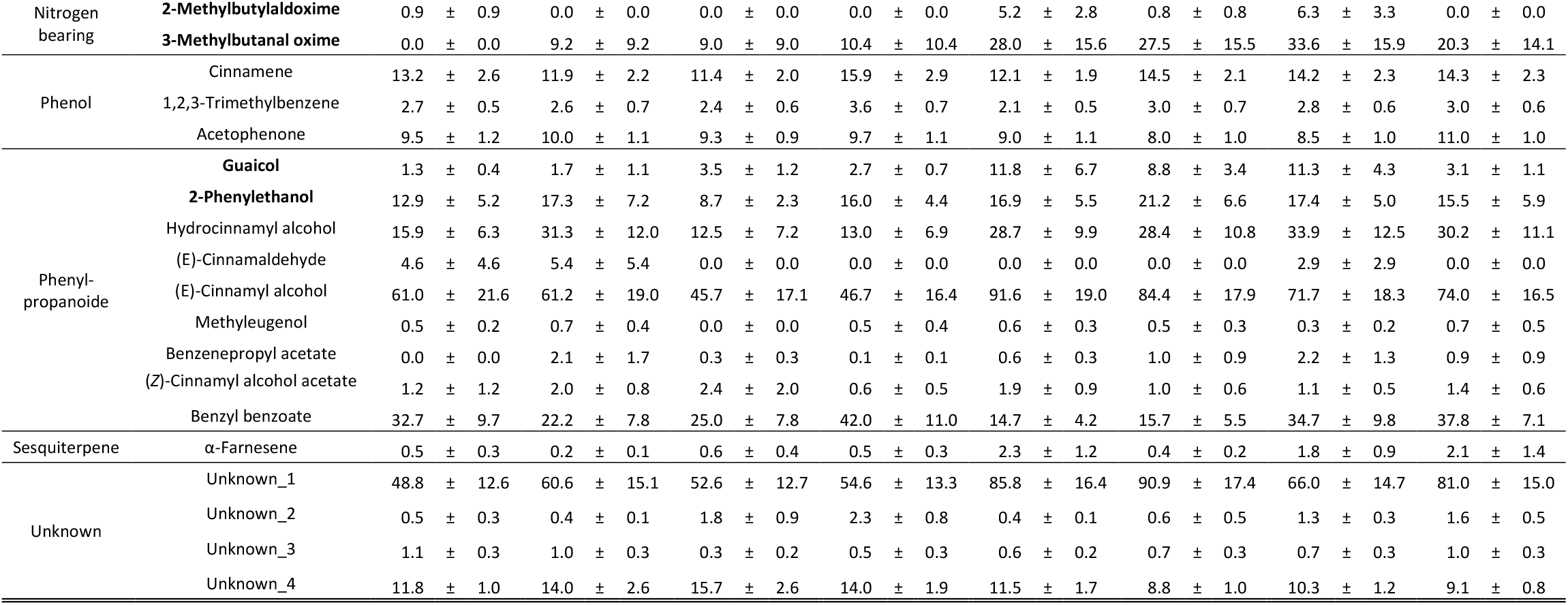
Mean ± SE abundance (e-06) of 70 volatile organic compounds (VOC) emitted by *Sielene latifolia* plants in all breeding treatment × sex × origin combinations, as determined by silicone tubing headspace collection combined with thermo desorption–gas chromatography–mass spectrometry. The 20 VOC, which evidently trigger antennal responses in *Hadena bicruris* (Dötterl et al., 2006) and were thus analysed as a sub-dataset, are highlighted in bold.

**Figure S6:**
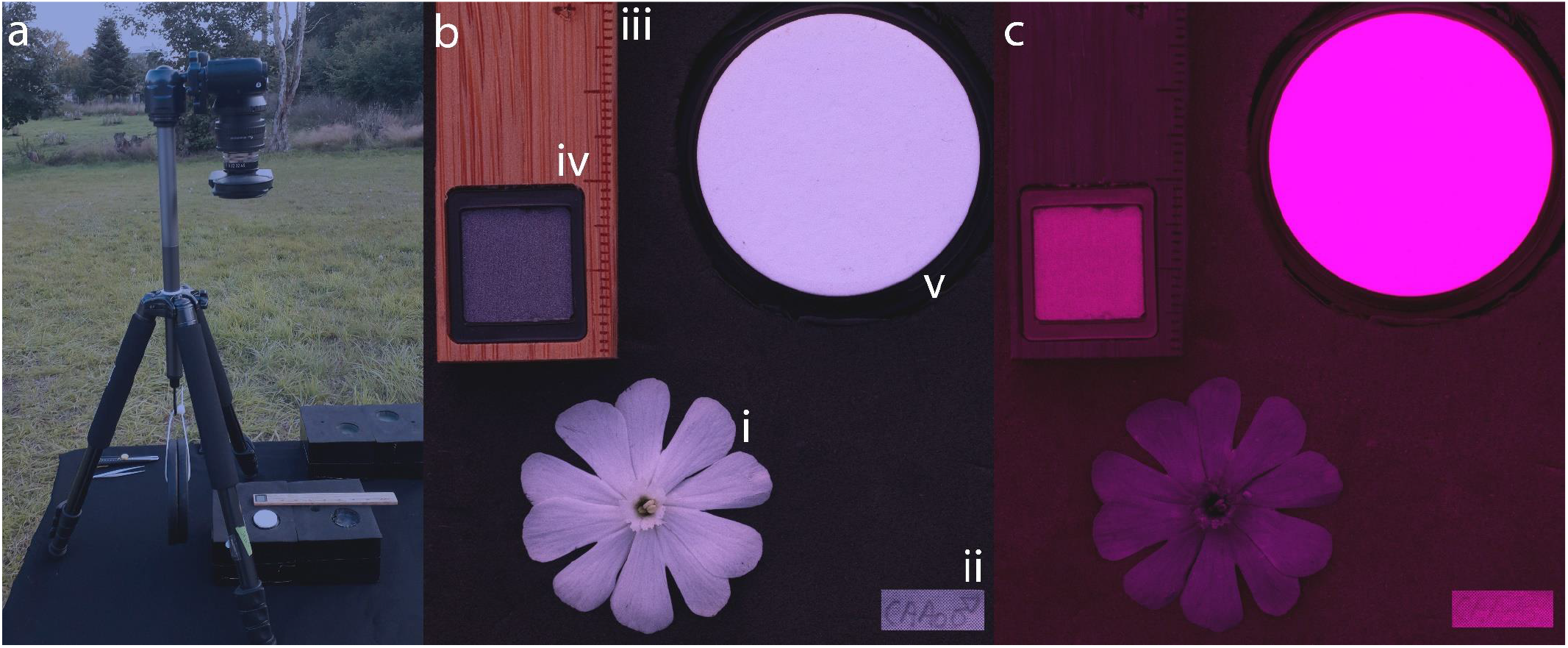
Setup for the acquisition of digital images for flower colour analyses. The camera was fixed on a tripod positioned on an exact horizontal platform, which was oriented towards the setting sun (a) to take images of flowers in the visible light spectrum (b) and the ultra violet light spectrum (c). Images included an intact and fully opened flower (i) that was carefully plugged into a black ethylene vinyl acetate sheet equipped with a label (ii), a size standard (iii), a 10 % polytetrafluorethylene light standard (iv) and a 99 % spectralon light standard (v).

**Figure S7:**
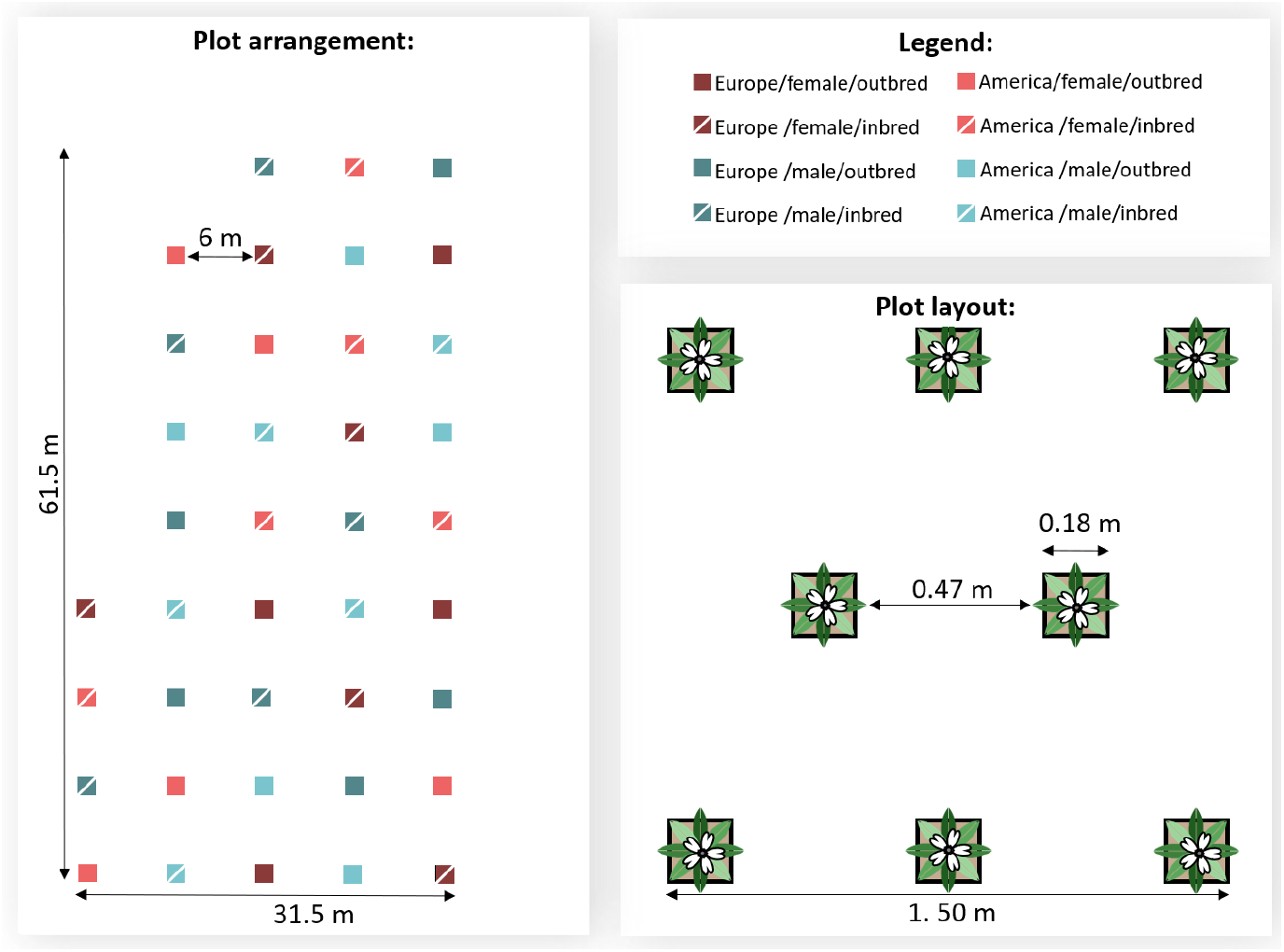
Experimental setup for pollinator observations. The plots consisted of eight individuals representing all eight populations within one of the eight possible breeding treatment × sex × range combinations. Each of these combinations was replicated five times on the maternal family level, resulting in 40 plots. Plots were spaced at a distance of 6 m to provide pollinators with the choice of visiting plants of specific breeding treatment × sex × range combinations.

### S8: Multivariate statistical analyses of VOC data

#### Methods S8a

Patterns of VOC abundance were compared in Random Forest (RF) models (R-packages: randomForest v4.6-14, party v1.3-5; Breiman, 2001). For each RF classification tree (the number of RF trees = iterations was set to 10,000), eight randomly selected variables were accepted as candidates at each split (mtry was set to 8, which is approx. the square root of the number of variables, i.e. the 70 compounds). The results from this unsupervised model are displayed in multi-dimensional scaling of proximity matrix (MDS) plots. The importance of each variable (i.e. chemical compound) in predicting the separation of data points was determined in separate supervised models and displayed as mean decrease in accuracy (MDA), if the variable was removed from the calculation. The approach was repeated for the 20 VOC, which evidently trigger antennal responses in *H. bicruris* (Dötterl et al., 2006), see Table S5.

**Figure S8b:**
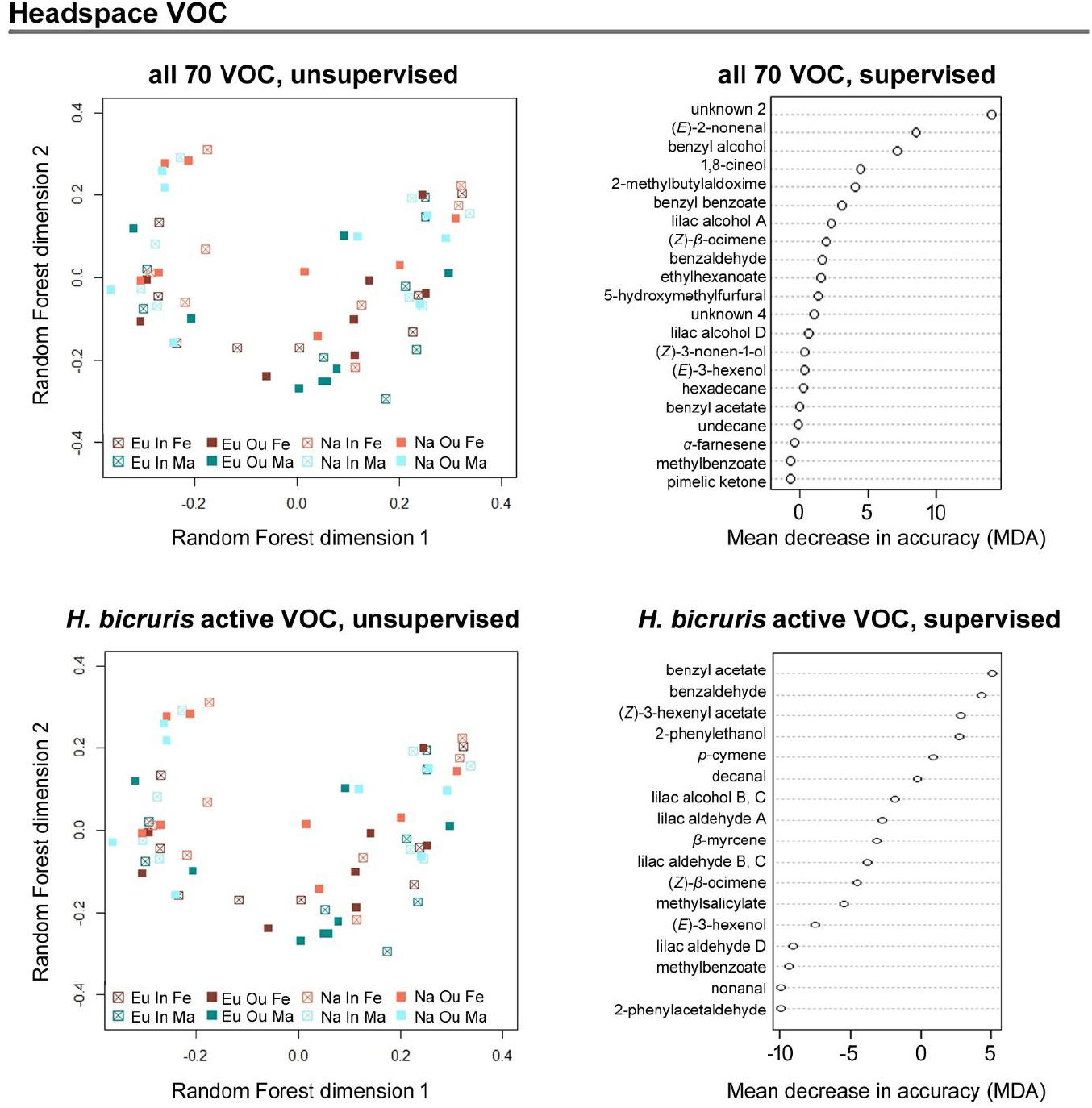
Unsupervised Random Forest comparison MDS plots for floral headspace VOC (left panel) and supervised Random Forest importance plots for mean decrease in accuracy (MDA, right panel) for all detected compounds (upper plots) and the subset for compounds that can be detected by *H. bicruris* (lower plots). Patterns were compared for outbred (Ou, filled squared) and inbred (In, open squares), female (Fe, red) and male (Ma, blue) plants from Europe (Eu, dark coloured) and North America (Na, bright coloured). Each square represents one population, data within populations were averaged to improve clarity

**Fig S9.**
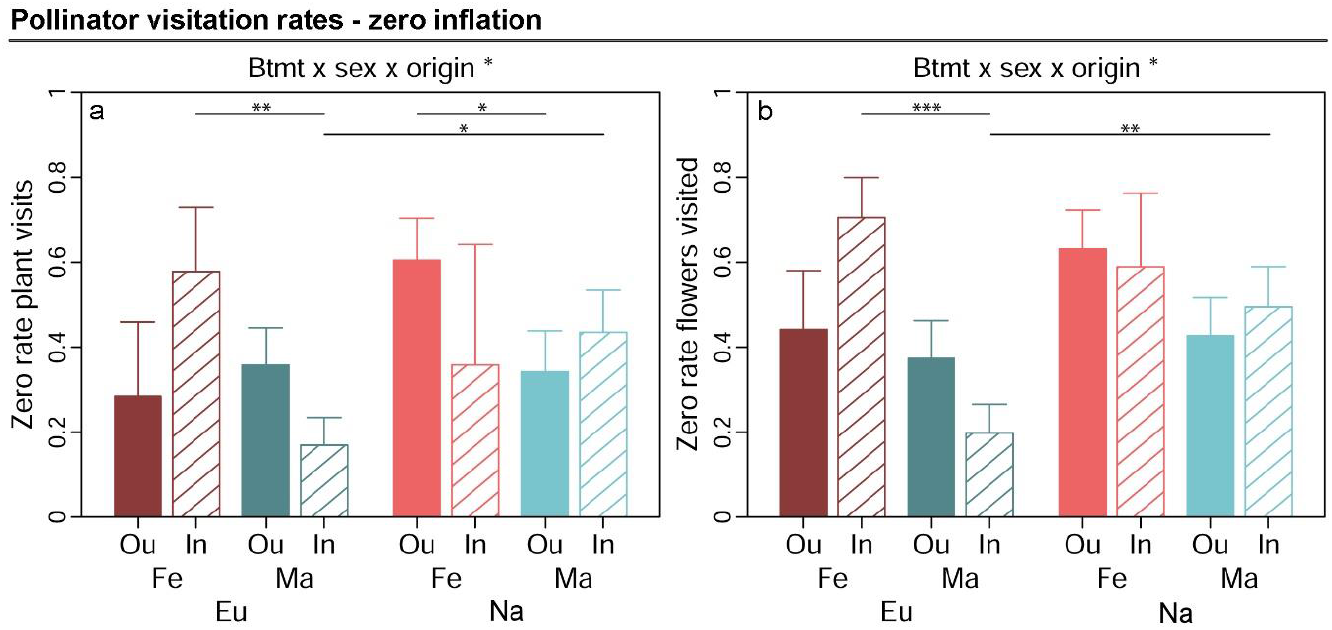
Graphs show estimated marginal means for zero scores in pollinator visitation responses (increasing values indicate higher proportion of zeroes in response data) and standard errors for outbred (Ou, filled bars) and inbred (In, open bars), female (Fe, red bars) and male (Ma, blue bars) plants from Europe (Eu, dark coloured bars) and North America (Us, bright coloured bars). Estimates were extracted for significant interaction effects from the zero-inflation part of generalised linear mixed effects models (significance levels based on Wald χ^2^-tests denoted at top of plot). Plots additionally indicate significant differences between breeding treatments, sexes or origins within levels of other factors involved in the respective interaction (estimated based on post-hoc comparisons, denoted within plots). Exact sample sizes for all responses are listed in Table 1. Significance levels: ***: p < 0.001, **: p < 0.01, and *p < 0.05.

The number of pollinator visits per plant was shaped by the interaction breeding treatment × sex × origin in the zero-inflation part of the model (p = 0.049, χ^2^_(1DF)_ = 4.30, Table 2, Fig. S9). Post-hoc comparisons revealed that zero-scores in plant visits were higher in females than males in European inbreds (p_post_ = 0.009) and North American outbreds (p_post_ = 0.037); and higher in North American than European plants for inbred males (p_post_ = 0.017). The number of flowers approached during a visit also depended on the interaction breeding treatment × sex × origin in the zero-inflation part of the model (p < 0.047, χ^2^_(1DF)_ = 4.15, Table 2, Fig. S9). Post-hoc comparisons revealed higher zero scores in females than males for European inbreds (p_post_ < 0.001) and higher zero scores in North American than European plants for inbred males (p_post_ = 0.006).

